# Developing an automated skeletal phenotyping pipeline to leverage biobank-level medical imaging databases

**DOI:** 10.1101/2022.10.31.514317

**Authors:** Chelsea C. Cataldo-Ramirez, David Haddad, Nina Amenta, Timothy D. Weaver

## Abstract

**Objectives:** Collecting skeletal measurements from medical imaging databases remains a tedious task, limiting the research utility of biobank-level data. Here we present an automated phenotyping pipeline for obtaining skeletal measurements from DXA scans and compare its performance to manually collected measurements.

**Materials and Methods:** A pipeline that extends and modifies the Advanced Normalization Tools (ANTs) framework was developed on 341 whole-body DXA scans of UK Biobank South Asian participants. A set of 10 measurements throughout the skeleton was automatically obtained via this process, and the performance of the method was tested on 20 additional DXA images by calculating percent error and concordance correlation coefficients (CCC) for manual and automated measurements. Stature was then regressed on the automated femoral and tibia lengths and compared to published stature regressions to further assess the reliability of the automated measurements.

**Results:** Based on percent error and CCC, the performance of the automated measurements falls into three categories: poor (sacral and acetabular breadths), variable (trunk length, upper thoracic breadth, and innominate height), and high (maximum pelvic aperture breadth, bi-iliac breadth, femoral maximum length, and tibia length). Stature regression plots indicate that the automated measurements reflect realistic body proportions and appear consistent with published data reflecting these relationships in South Asian populations.

**Discussion:** Based on the performance of this pipeline, a subset of measurements can be reliably extracted from DXA scans, greatly expanding the utility of biobank-level data for biological anthropologists and medical researchers.

## Introduction

The advent of large medical imaging databases has the potential to expand the bounds of research for many disciplines, including anthropology. This is particularly true for investigations into the relationships between genotype and phenotype, which necessitate very large samples to offset power limitations incurred by the size of the human genome (Asif et al., 2021). A better understanding of genotype-phenotype relationships in humans could be transformative for many areas of biological anthropology, including testing evolutionary hypotheses about human biological variation or the fossil record for human evolution. Extracting relevant phenotypic data from the large biobanks containing matched genotypic data, however, remains tedious and in certain contexts, effectively infeasible because of the time required.

To obtain quantitative skeletal information from imaging databases, researchers are typically limited to two options: manually collecting the data, or implementing an automated procedure. For the manual option, the images are visually assessed and interpreted by an experienced researcher. This approach usually results in specific, detailed, and reliable data, but the process is typically time intensive, which limits the number of images that can be included in the study, particularly when the data is only available to the researcher for a limited time frame. Automated methods of phenotyping have the potential to greatly increase the number of images that can be studied as they can extract much more data in a shorter period of time. This benefit is potentially offset, however, by the quality and reliability of the data. In an effort to bridge the gap between the quality versus quantity tradeoff for obtaining quantitative data from medical images, we assess the performance of an existing image registration framework (Advanced Normalization Tools [ANTs]) on automating the extraction of postcranial skeletal measurements from total body dual-energy x-ray absorptiometry (DXA) scans. Here, we develop a pipeline to process DXA images, extract biologically meaningful skeletal measurements, and optimize the accuracy and reliability of the auto-generated measurements on a small subset of UK Biobank participants, which can then be applied to larger datasets that cannot reasonably be phenotyped by manual methods.

## Materials & Methods

### Sample

The data used for this study consists of a small subset of whole-body DXA images available from the UK Biobank (UKB), which contains demographic, lifestyle, anthropometric, health-related, genetic, and imaging data for over 500,000 adult individuals residing in the United Kingdom (Bycroft et al., 2018; Littlejohns et al., 2020). Whole body DXA images provide an anterior-posterior view of the entire body, with enough resolution to extract body composition measures such as bone mineral density as well as 2D linear measurements of skeletal features. DXA scanning requires a much lower dose of radiation than X-rays, making them a safer imaging method for patients, although at the cost of losing some finer-tuned resolution (Njeh et al., 1997). While there are approximately 47,000 participants with whole body DXA images (https://biobank.ndph.ox.ac.uk/showcase/field.cgi?id=20158), our sample is composed solely of participants who self-identify as one of three South Asian groups: Indian, Pakistani, or Bangladeshi (Table 1). This sub-sample was selected in an effort to move away from the Eurocentric bias in biobank-level research and shift the focus towards groups that are typically underrepresented in this domain (e.g., Extended Data Table 3 of Bycroft et al., 2018) and to develop the automated phenotyping pipeline on a sample that was small enough to be investigated in detail. Limiting the study to this sub-sample resulted in a significant reduction in the number of available whole body images (*n* = 361 after quality control assessments). The 361 images were divided into “atlas”, “training”, and “test” samples for the study (Figure 1; Table 1). Thirteen male and female (*n* = 26) atlas images were selected to act as templates for guiding the deformation and landmark propagation process (described in more detail in the *Methods* section) and ten randomly selected male and female (*n* = 20) images were set aside for error assessments. The remaining 315 images comprised the training sample, upon which the parameters and procedures described below were refined. The data utilized in this study fall generally under the ethics approvals granted to the UKB. The UKB received approval from the National Information Governance Board for Health and Social Care and the National Health Service North West Centre for Research Ethics Committee (Ref: 11/NW/0382), maintains its own internal ethics board (UK Biobank Ethics Advisory Committee), and is compliant with the General Data Protection Regulation (GDPR) (Bycroft et al., 2018; Littlejohns et al., 2020). Participants enrolled in the UKB provided electronic signed consent at the time of enrollment (Bycroft et al., 2018; Littlejohns et al., 2020).

**Table 1:** Summary of the partitioning to the total sample. In total, 361 images were used: 315 in the training set, 26 as atlases, and 20 were assigned to the test set.

|  | Atlas |  | Training |  | Test |  | Total |
| --- | --- | --- | --- | --- | --- | --- | --- |
|  | *M* | *F* | *M* | *F* | *M* | *F* |  |
| **Indian** | 10 | 9 | 144 | 80 | 6 | 6 | 255 |
| **Pakistani** | 3 | 4 | 45 | 25 | 2 | 2 | 81 |
| **Bangladeshi** | 0 | 0 | 15 | 6 | 2 | 2 | 25 |
| **Total** | 13 | 13 | 204 | 111 | 10 | 10 | 361 |

**Table 2:** Measurements and landmarks used in this study. Extended definitions are available in SI Table 1.

| Measurement | Landmarks |  |
| --- | --- | --- |
| Trunk length | T1 | S1 |
| Upper thoracic breadth | R4L | R4R |
| Superior sacral breadth | SSIL | SSIR |
| Inferior sacral breadth | ISIL | ISIR |
| Innominate height | InSL/R | IsIL/R |
| Bi-iliac breadth | BIBL | BIBR |
| Maximum pelvic aperture | PAL | PAR |
| Acetabular breadth | AcSL/R | AcIL/R |
| Femoral maximum length | FSL/R | FDL/R |
| Tibia length | TPL/R | TDL/R |

**Table 3:**
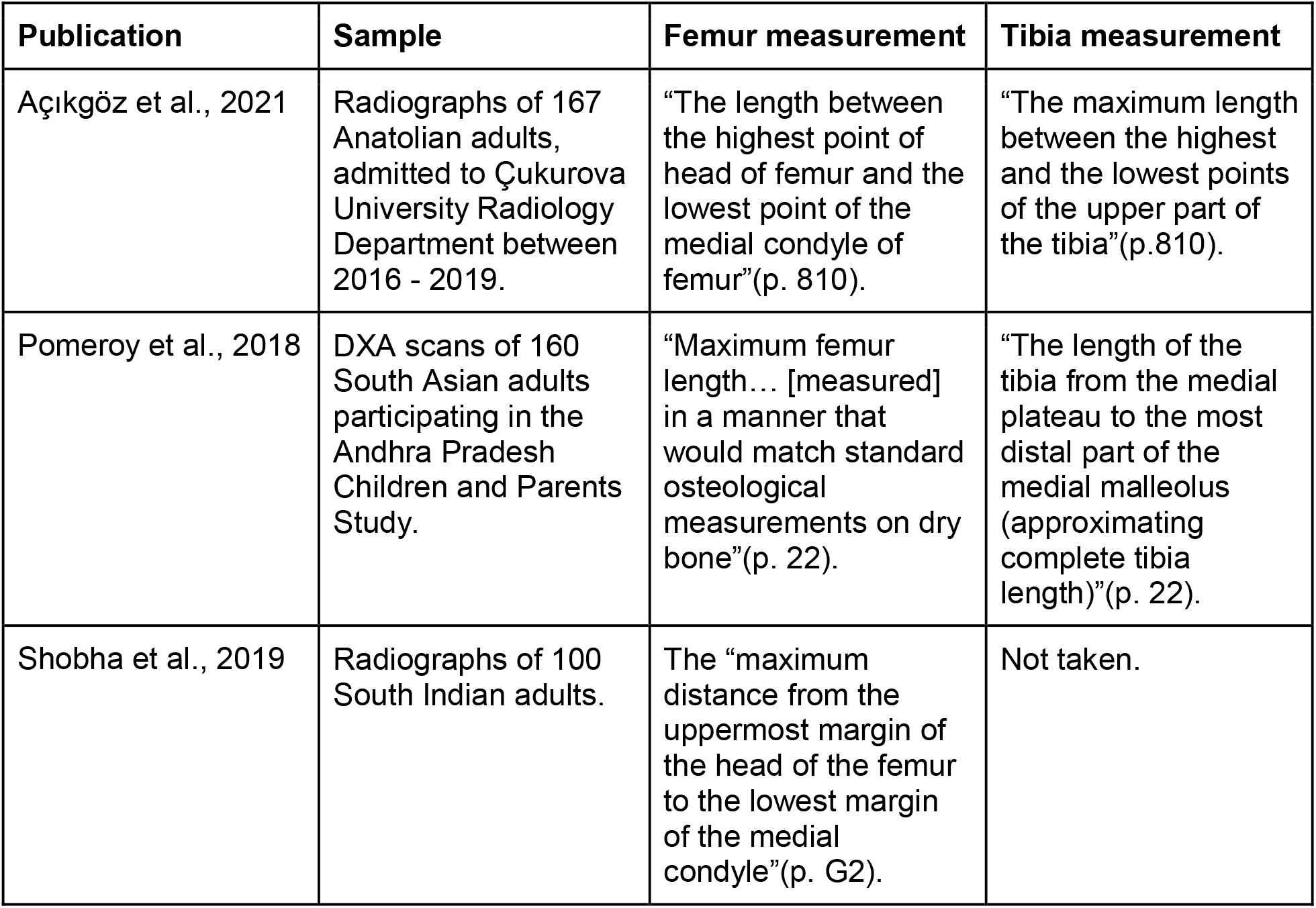
Sample information and measurement definitions for publications used to compare stature regression equations incorporating lower limb lengths. It is important to note that the two tibia measurements differ from each other and from the tibia length used in this study. Unlike the measurement used in this study, Pomeroy et al. incorporated the medial malleolus in their measurement, which more closely resembles how one would take a maximum tibia length on dry bone. It is unclear exactly how Açlkgöz et al. define tibia length based on their definition alone, and a figure of the measurement was not provided.

**Figure 1:**
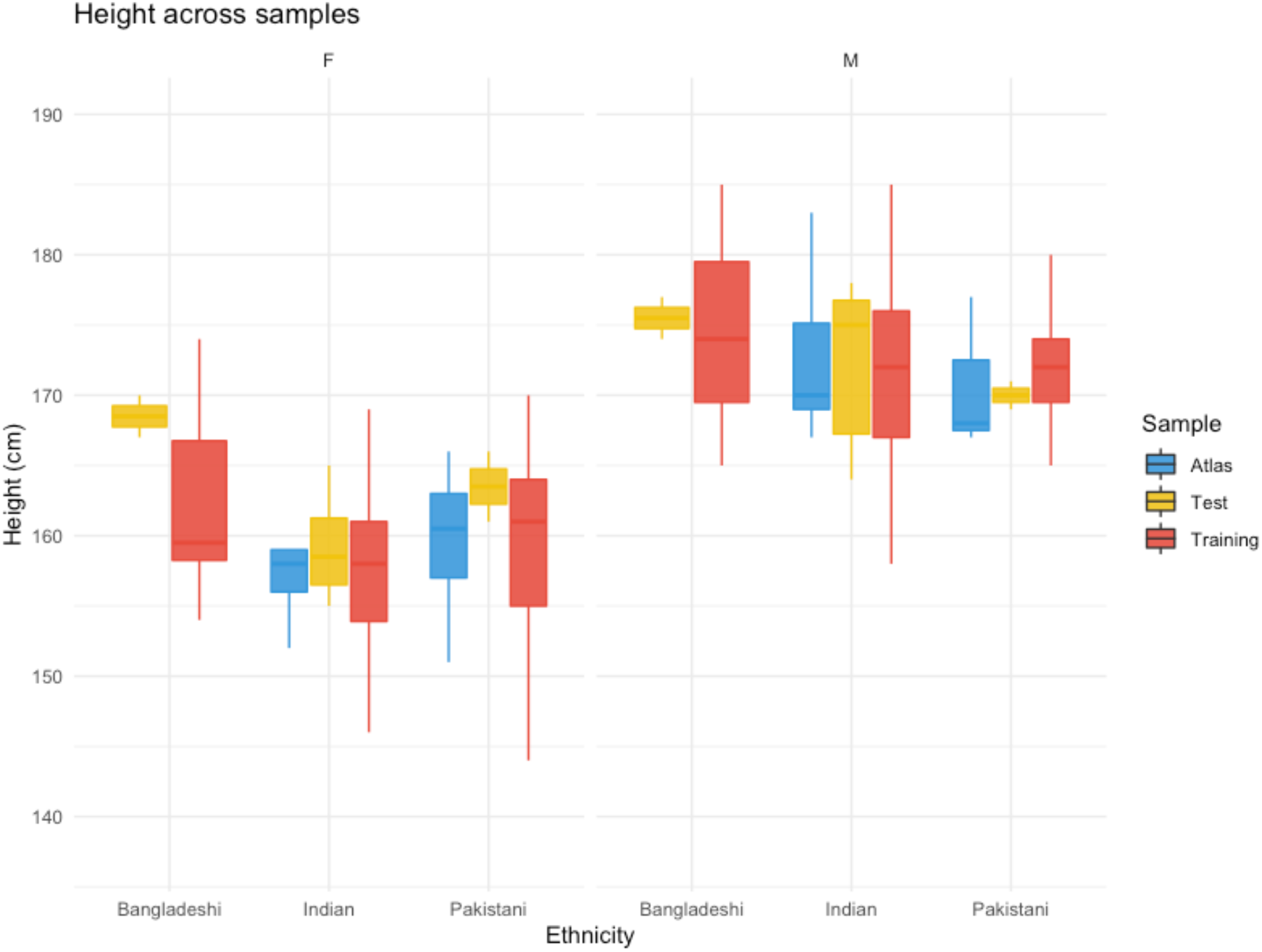
UKB reported height (UKB data field 50-0.0) for participants whose DXA images were used in this study, grouped by sex, self-reported ethnicity, and sample.

### Methods

#### Advanced Normalization Tools (ANTs)

The underlying methodology for our automated phenotyping procedure relies on the ANTs framework, which was originally derived from the Insight Toolkit, used for analyzing 3D medical images of brains (Avants, Tustison, and Song, 2009; Avants, Johnson, and Tustison, 2015). ANTs can be used to automate landmark propagation between an atlas (or “fixed”) image and any number of other images (“moving” images) of the same type, vastly minimizing the time needed for manual landmark placement (Avants, Tustison, and Johnson, 2014). Briefly, our use of ANTs consists of manually landmarking an atlas, registering and warping each moving image to the atlas, and using inverse warping to propagate the landmarks from the atlas to the moving image (Figure 2). The registration and warping step consists of a rigid transformation to align the moving to the atlas image, an affine transformation to match the moving to the atlas image by scaling and shearing, and a non-linear transformation to match localized regions of the moving to the atlas image. Before adding to the pipeline, the registration framework was optimized for use with 2D whole body DXA scans by experimenting with parameter modifications and visualizing the resulting landmark propagations. When visual inspections no longer yielded clear improvements in skeletal landmark propagation, we formally tested the performance of ANTs on automating the extraction of postcranial skeletal measurements on a small subset of DXA images from the UKB but not part of the South Asian sub-sample (Cataldo-Ramirez et al., 2022). The results of this preliminary investigation showed that using ANTs registration and landmark propagation alone yielded high variance in percent error across all skeletal measurements assessed, and that not all atlases perform equally well for any given moving image. This highlighted several limitations as well as areas to prioritize for further optimization. Image resolution of the DXA scans and (non-standardized) limb positioning of the participants constrain which skeletal measurements can realistically be automatically extracted with a pipeline based on ANTs, leading to a reduction in skeletal metrics tested here (Figure 3, Table 2, SI Table 1). Additionally, the variance in performance across atlas images suggests that significant consideration should be given to selecting which DXA images should be used as atlases, and that using more than one atlas could increase performance. These insights guided the development of the protocols implemented for the current study.

**Figure 2:**
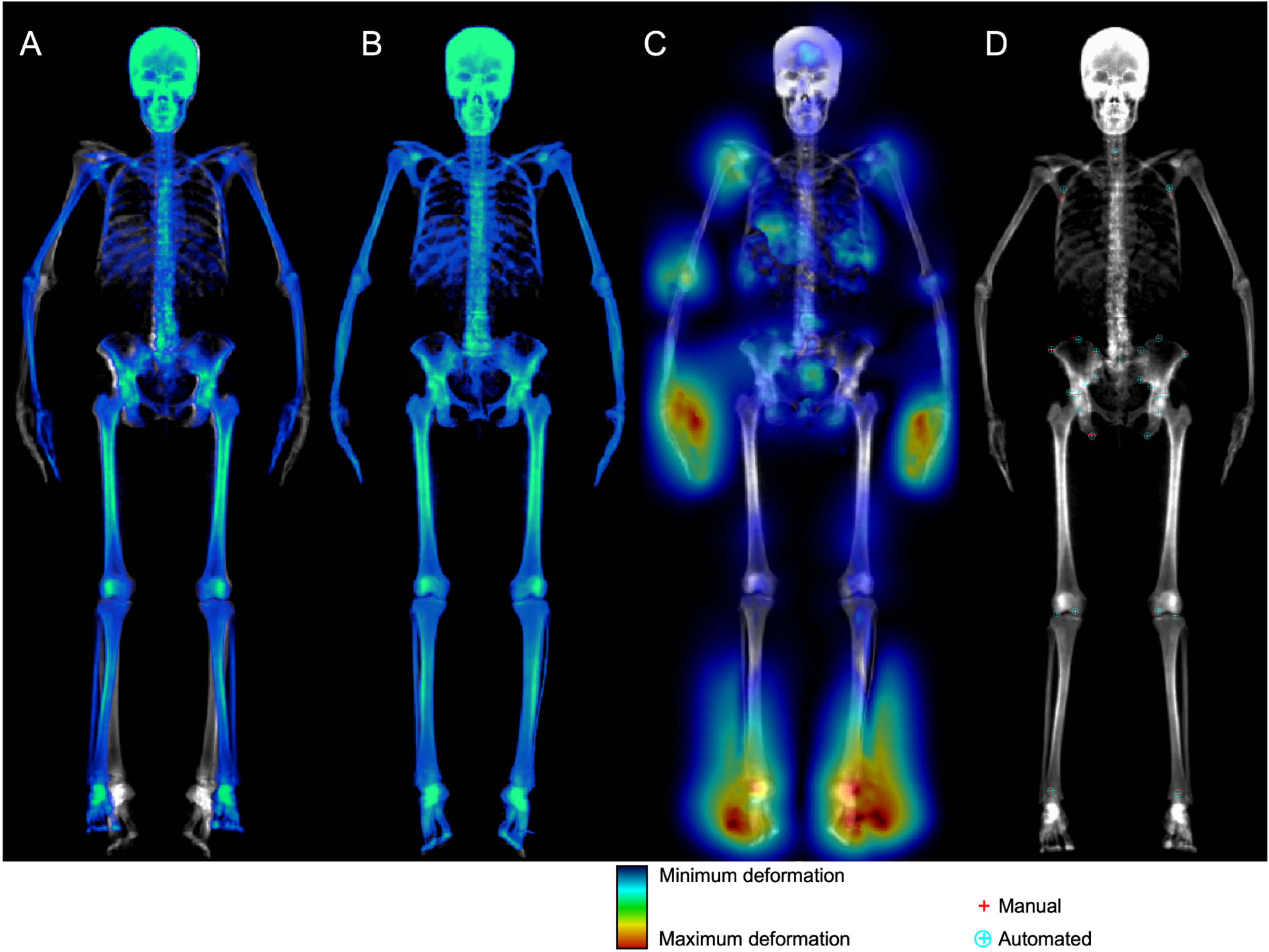
Image registration and landmark propagation between an atlas image and a moving image. A) The combined rigid and affine transformation of the moving image (color) overlaid on the atlas (black and white); B) the final, non-linear transformation of the moving image overlaid on the atlas; C) the inverse-transformation (color) highlighting the location and magnitude of deformations needed to adjust the propagated landmarks from the warped moving image (black and white) back to its original proportions; D) the moving image with both manual (red plus signs) and propagated (cyan circles with crosshair) landmarks plotted. Images reproduced by kind permission of UK Biobank ©.

**Figure 3:**
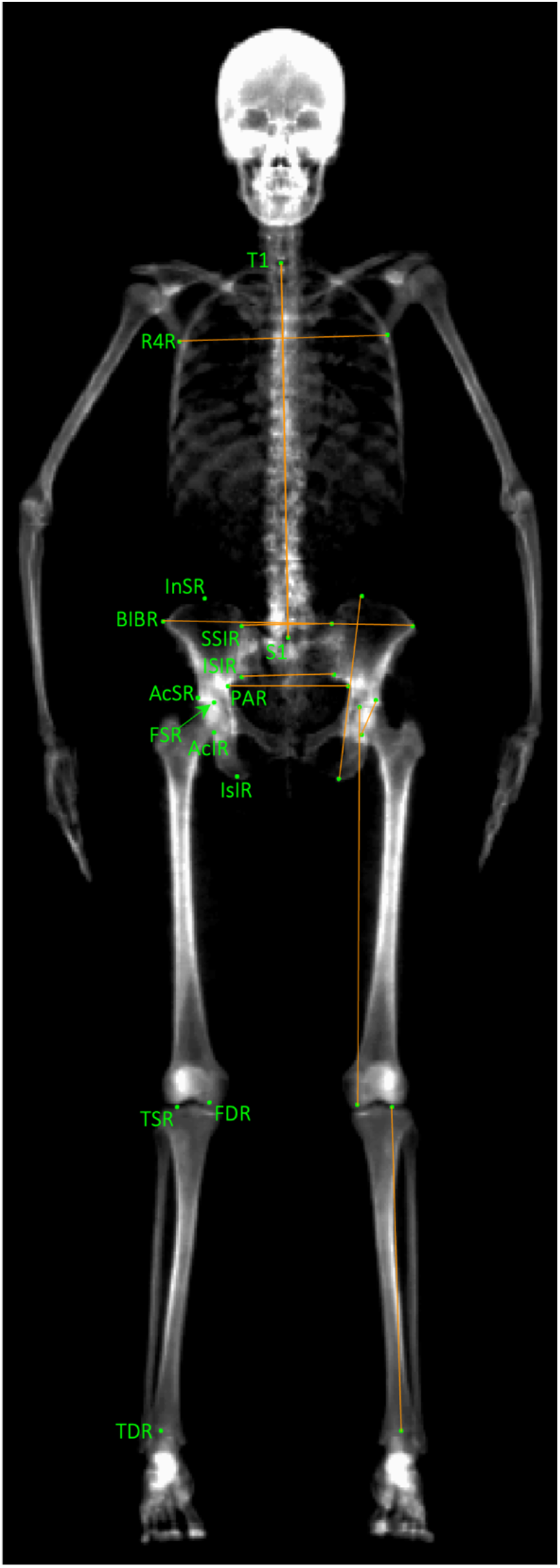
An example atlas, or “fixed image”, with manually placed landmarks (green circles). Landmark definitions were constrained by image quality and 2D compression of 3D anatomy. Labels (green text) are provided only on the right side for bi-lateral landmarks. Inter-landmark distances (orange lines) are overlaid only on the left side for bi-lateral metrics. Images reproduced by kind permission of UK Biobank ©.

#### Image Pre-processing

The DXA scans, as provided by the UKB, do not follow strict standardization in image processing and therefore were subjected to a pre-processing pipeline to optimize image uniformity. This included converting the images from DICOM to NIftI format (preferred by ANTs), converting all image backgrounds to black, adjusting the contrast to best visualize skeletal elements and remove soft tissue, and outputting a grid figure of all processed images for manual quality control inspection (Figure 4). All images were pre-processed using MATLAB R2022a and the dicm2nii add-on (Mathworks, 2022; Li, X., 2022).

**Figure 4:**
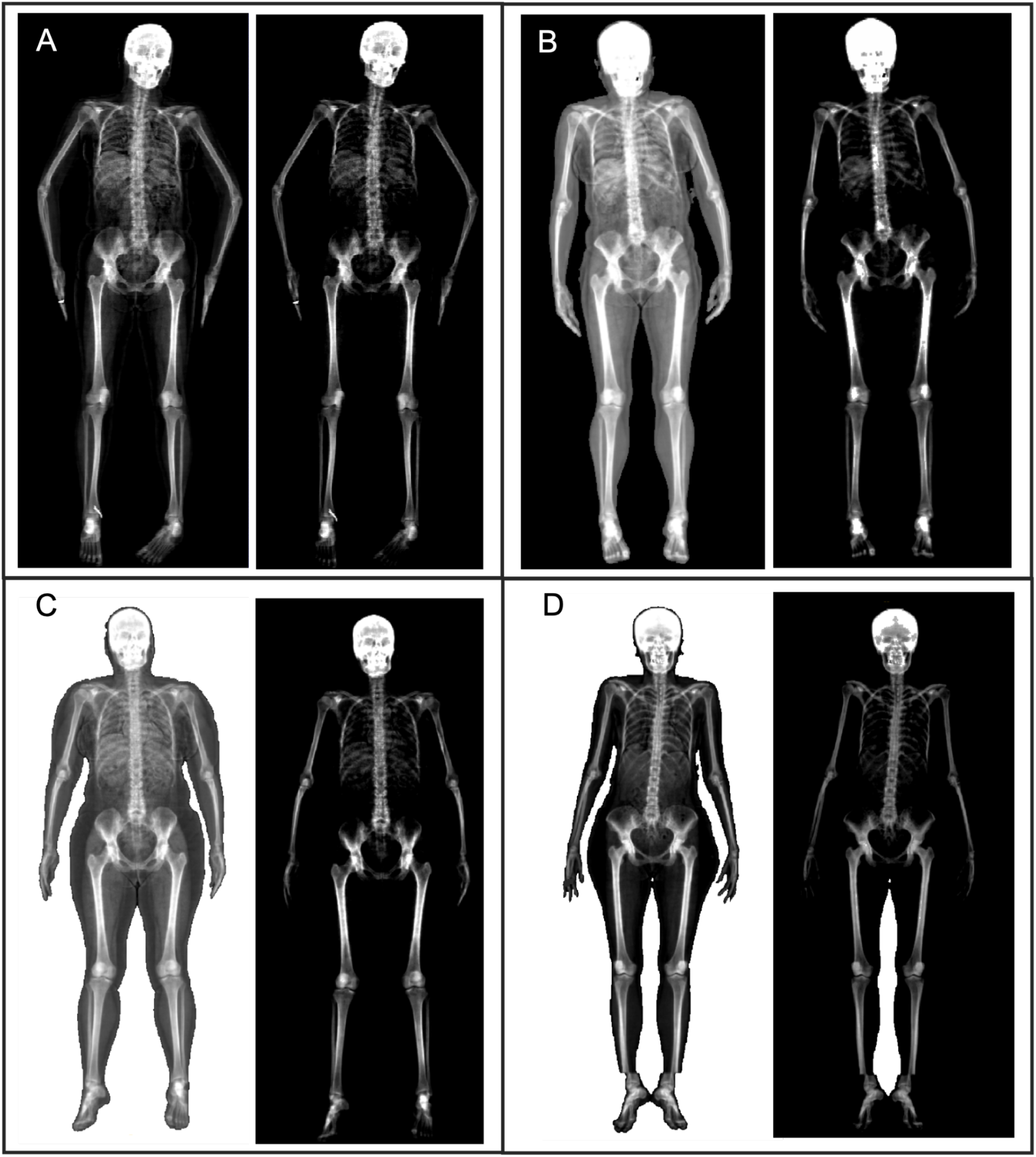
DXA scans of four individuals as received by the UKB (left) and after automated image preprocessing for uniformity (right). The left panels of A-D demonstrate some of the variation in body positioning, image contrast, and background color observed within the sample. D was removed during the visual inspection of the grid figure output by the image preprocessing pipeline due to the imaging artifact just proximal to the ankles. D also highlights the limitation of the current image preprocessing pipeline at homogenizing the background color in images with white backgrounds and limbs touching, as this creates pockets or “islands” of white between the limbs. Images with identified white “islands” were manually processed to ensure their uniformity with the rest of the sample. Reviewing the grid image output after preprocessing is highly recommended for identifying scans unsuitable for automated phenotyping. Images reproduced by kind permission of UK Biobank ©.

#### Atlas Selection Procedure

After processing the images for uniformity, the scans were separated by self-reported sex (UKB data field 31-0.0). Within each group (males and females), elliptical Fourier analysis (EFA) was applied to the whole body outline, and principal component analysis was conducted on the resulting EFA harmonic coefficients. Hierarchical clustering using Ward’s criterion was then performed on the first three principal components. Five and three cluster solutions were suggested for males and females, respectively. The images closest to each cluster center were initially chosen as atlases. However, after reviewing the performance of subsequent ANTs runs on the training sample we incrementally increased the number of atlases. Ultimately, images representing both the cluster centers and cluster edges (i.e., images with the furthest distances from cluster centers) were incorporated, resulting in a total of 13 atlases per sex group. Atlas selection analyses were conducted using R Statistical Software (v4.2.0; R Core Team 2022) packages FactoMineR (Lê et al., 2008) and factoextra (Kassambara, A., Mundt, F., 2020) and MATLAB base functions and add-on “Elliptic fourier for shape analysis” (Mathworks, 2022; Manurung, A., 2022). After selecting the atlases, each image was manually landmarked (Figure 3; Table 2) in ITK-Snap (v3.6.0; Yushkevich et al., 2006) and the landmark coordinates (in pixels) were exported.

#### Post-propagation Quality Control

The ANTs registration and landmark propagation process was performed once for each atlas on all training images of the same sex, outputting 13 landmark sets per training image. To reduce error and lessen the likelihood of obtaining inaccurate landmark propagations, we employed a post-propagation outlier identification, landmark consistency check, and landmark averaging step. The goal of the outlier identification step is to detect cases where the transformations do not reflect anatomy, usually because of a poor match between the atlas and moving image. The goal of the consistency check is to detect inconsistencies among different propagations. The goal of the landmark averaging procedure is to statistically reduce error by basing the final landmark positions on multiple propagations. For the outlier identification step, first, the atlas landmark sets of both sexes were pooled and the Procrustes mean shape was calculated. Next, the Procrustes distances from the atlas mean to all 13 propagated landmark configurations per training image were calculated. The Procrustes mean shape and distances are calculated after the landmark configurations have been superimposed to remove the size, location, and rotation of the configurations (Dryden and Mardia, 1998). Importantly, because size has been removed, outliers are not those images that are particularly small or large; they are those that have unusual relationships among the landmarks. Propagated configurations with Procrustes distances exceeding a threshold that was empirically determined (tuned) on the training sample were excluded from further analyses. For the landmark consistency check, first, random triplets of all remaining configurations (per training image) were selected, and the average Euclidean distance between all matched landmark pairs in the triplet was calculated. Next, triplets with an average Euclidean distance below an empirically determined threshold were retained. Euclidean distance is used because each configuration in the triplet should represent the locations of landmarks on the same moving image, so size and location (as well as shape) are important for assessing the concordance of the configurations. For the landmark averaging step, each landmark in the accepted triplet set was then averaged and scaled from pixels (pix) to millimeters (mm). Lastly, the skeletal measurements were calculated from the landmark coordinates. Any training image that failed to pass through the landmark averaging quality checks was flagged for manual visual inspection, and no associated skeletal measurements were output.

#### Training Sample

All of the procedures described above were implemented on the training sample, which consisted of all images not used as atlases or in the error assessment analyses (the test sample). These images were used to refine the pipeline by visualizing the propagated landmarks on their corresponding moving images after every pipeline adjustment. The pipeline procedures and thresholds were finalized once we received a 90% acceptance rate from the post-propagation quality control process.

#### Error Assessment

The test sample (Table 1), composed of 10 randomly selected male and female (*n* = 20) images, was set aside after image pre-processing and before the atlas selection procedure. These images were manually landmarked and subjected to the finalized automated phenotyping pipeline. To assess the performance of the pipeline, the absolute differences between the automated and manually obtained skeletal measurements of the test sample images were calculated and then divided by the manually-derived skeletal measurements to obtain percent errors. A bi-variate plot of the manual and automated measurements that included the individual data points, a 1:1 line, and a least-squares regression of the manual on the automated measurement was used to visualize the bias and correlation between the automated and manual measurements. Lin’s concordance correlation coefficient (CCC) was also calculated for each measurement to quantify the deviation from a 1:1 association. All error analyses were conducted using R Statistical Software (v4.2.0; R Core Team 2022).

#### Comparisons with Stature and Other Studies

To further ensure the validity of the automated skeletal measurements, all measurements were plotted against stature (UKB data field 50-0.0). Stature was manually recorded by the UKB independent of the DXA image, so it therefore acts as an external proxy to assess if the skeletal measurements derived from the scans are realistic. Additionally, relationships between lower limb lengths and stature are well characterized and can provide a frame of reference for assessing the utility and validity of our automated skeletal measurements. Therefore, univariate regression equations for stature were calculated using maximum femur length and tibia length and were visually compared with published stature regression equations from Açikgöz et al., 2021, Pomeroy et al., 2018, and Shobha et al., 2019 (Table 3). The aforementioned publications present univariate stature regression equations derived from skeletal measurements taken from antero-posterior (AP) DXA scans or radiographic images from contemporary populations in or near South Asia. Relevant details of these publications are noted in Table 3.

#### Code Repositories / Data Availability

The scripts developed for the pipeline presented in this study will be made available upon request to the corresponding author. The skeletal measurements will be added to the UKB database and made available to researchers approved by the UKB review panel.

## Results

Of the 20 images in the test sample, one male and one female image failed the post-propagation outlier identification and landmark averaging procedure. This resulted in a 90% acceptance rate, mirroring the acceptance rate of the training sample. Percent errors for the skeletal measurements of the passing images ranged from 0.04% (average femoral maximum length) to 29.97% (inferior sacral breadth) (Table 4). Minimum CCC values at or above 0.90 (indicating acceptable performance [Altman, 1991]) were achieved for bi-iliac breadth (95% CI = 0.92 - 0.99), max pelvic aperture (95% CI = 0.93 - 0.99), average max femoral length (95% CI = 0.93 - 0.99), and average tibia length (95% CI = 0.90 - 0.98) (Table 5). Based on the percent error and CCC, the performance of the automated skeletal measurements can be characterized into three categories: poor (sacral and acetabular breadths), variable (trunk length, upper thoracic breadth, and innominate height), and high (max pelvic aperture breadth, bi-iliac breadth, femoral max length, and tibia length). The association between the manual and automated measurements and the bias of the automated measurements is visualized in Figure 5. Only bi-iliac breadth appears to be minimally biased (the automated measurement is underestimated by 2.23 mm on average) among the skeletal measurements with acceptable CCC values (Fig. 5C).

**Table 4:** Percent errors between manual and automated skeletal measurements in the test sample. Percent error was calculated as the absolute value of the difference between manual and automated measurement divided by the manual measurement.

| Measurement | Min (%) | Max (%) | Mean (%) |
| --- | --- | --- | --- |
| Trunk length | 0.38 | 6.18 | 2.22 |
| Upper thoracic breadth | 0.20 | 6.18 | 2.51 |
| Superior sacral breadth | 0.91 | 17.92 | 8.41 |
| Inferior sacral breadth | 0.16 | 29.97 | 9.28 |
| Innominate height* | 0.07 | 6.87 | 2.79 |
| Bi-iliac breadth | 0.10 | 2.06 | 1.04 |
| Maximum pelvic aperture | 0.06 | 3.47 | 1.28 |
| Acetabular breadth* | 1.32 | 18.73 | 6.38 |
| Femoral maximum length* | 0.04 | 3.49 | 0.70 |
| Tibia length* | 0.06 | 3.66 | 1.13 |

\*The average for bilateral metrics is reported.

**Table 5:** Lin’s concordance correlation coefficient (CCC) for manually-derived and automated measurements (test sample).

| **Measurement** | **CCC** | **Lower 95% CI CCC** | **Upper 95% CI CCC** |
| --- | --- | --- | --- |
| Trunk length | 0.85 | 0.64 | 0.94 |
| Upper thoracic breadth | 0.90 | 0.77 | 0.96 |
| Superior sacral breadth | 0.44 | 0.02 | 0.73 |
| Inferior sacral breadth | -0.11 | -0.42 | 0.23 |
| Innominate height* | 0.81 | 0.57 | 0.92 |
| Bi-iliac breadth | 0.97 | 0.92 | 0.99 |
| Maximum pelvic aperture | 0.97 | 0.93 | 0.99 |
| Acetabular breadth* | 0.35 | -0.03 | 0.64 |
| Femoral maximum length* | 0.97 | 0.93 | 0.99 |
| Tibia length* | 0.96 | 0.90 | 0.98 |

\*The average for bilateral metrics is reported.

**Figure 5:**
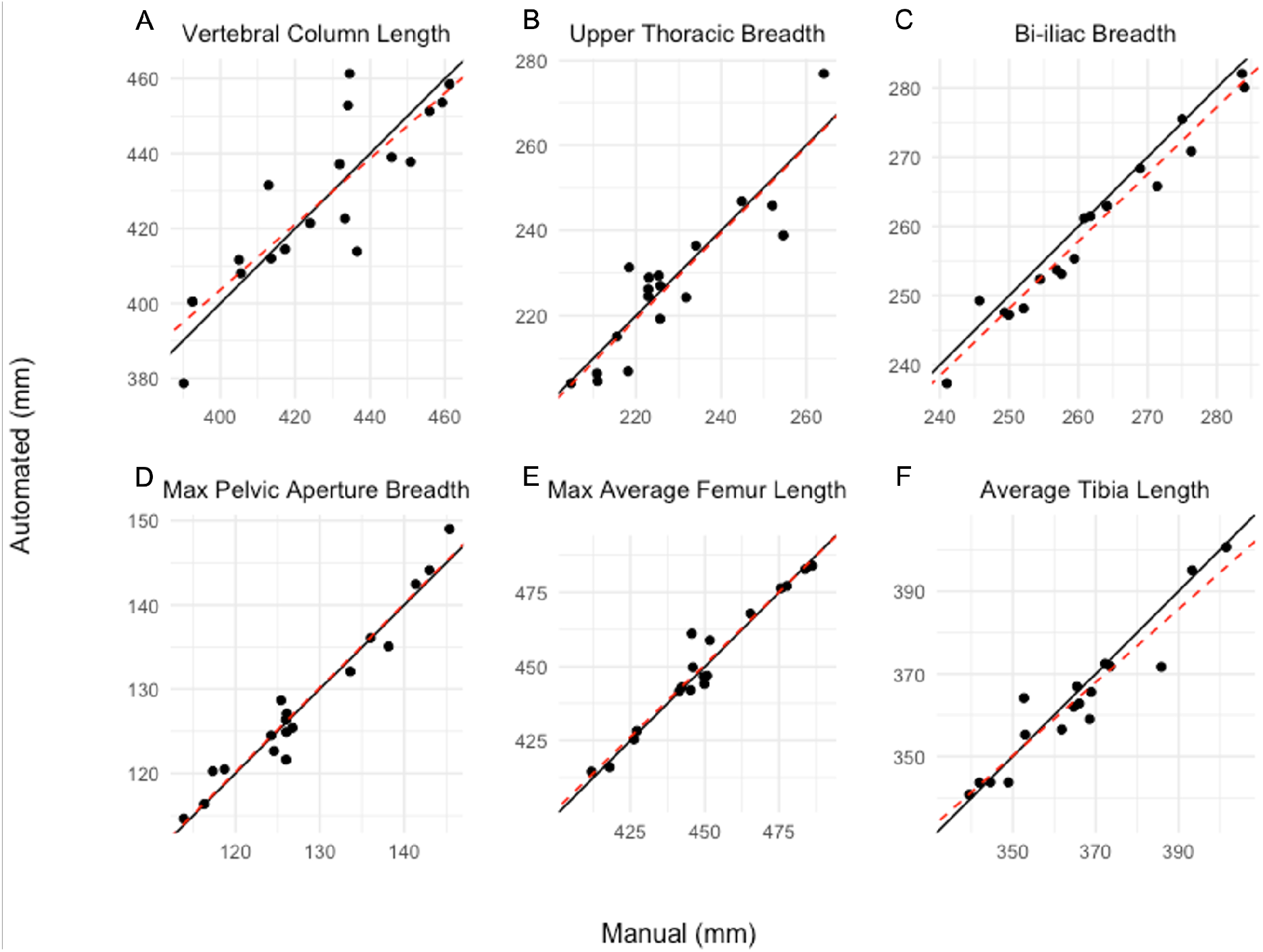
A-F) Association between manually-derived (x-axis) and automated (y-axis) skeletal measurements of the test sample. The solid black lines represent a 1:1 concordance as a reference. The black circles represent the individual data points and the dashed red lines represent the least-squares regressions of the automated on the manual-derived measurements.

Stature regression plots (Figure 6) indicate that the automated skeletal measurements reflect realistic body proportions, and that there is reasonable concordance between manually recorded stature and 2D image-derived measurements. Comparisons with previously published data reflecting the relationships between stature and lower limb lengths in South Asian populations appear consistent with those presented here (Açikgöz et al., 2021, Pomeroy et al., 2018; Shobha et al., 2019) (Table 3, Figure 6). With the exception of Pomeroy et al.’s tibia equation, there is considerable overlap between published regression lines and our data points (Figure 6). However, Pomeroy’s tibia measurement includes the medial malleolus while ours does not, therefore increasing their tibia measurement and likely explaining the discrepancy between our stature regression lines.

**Figure 6:**
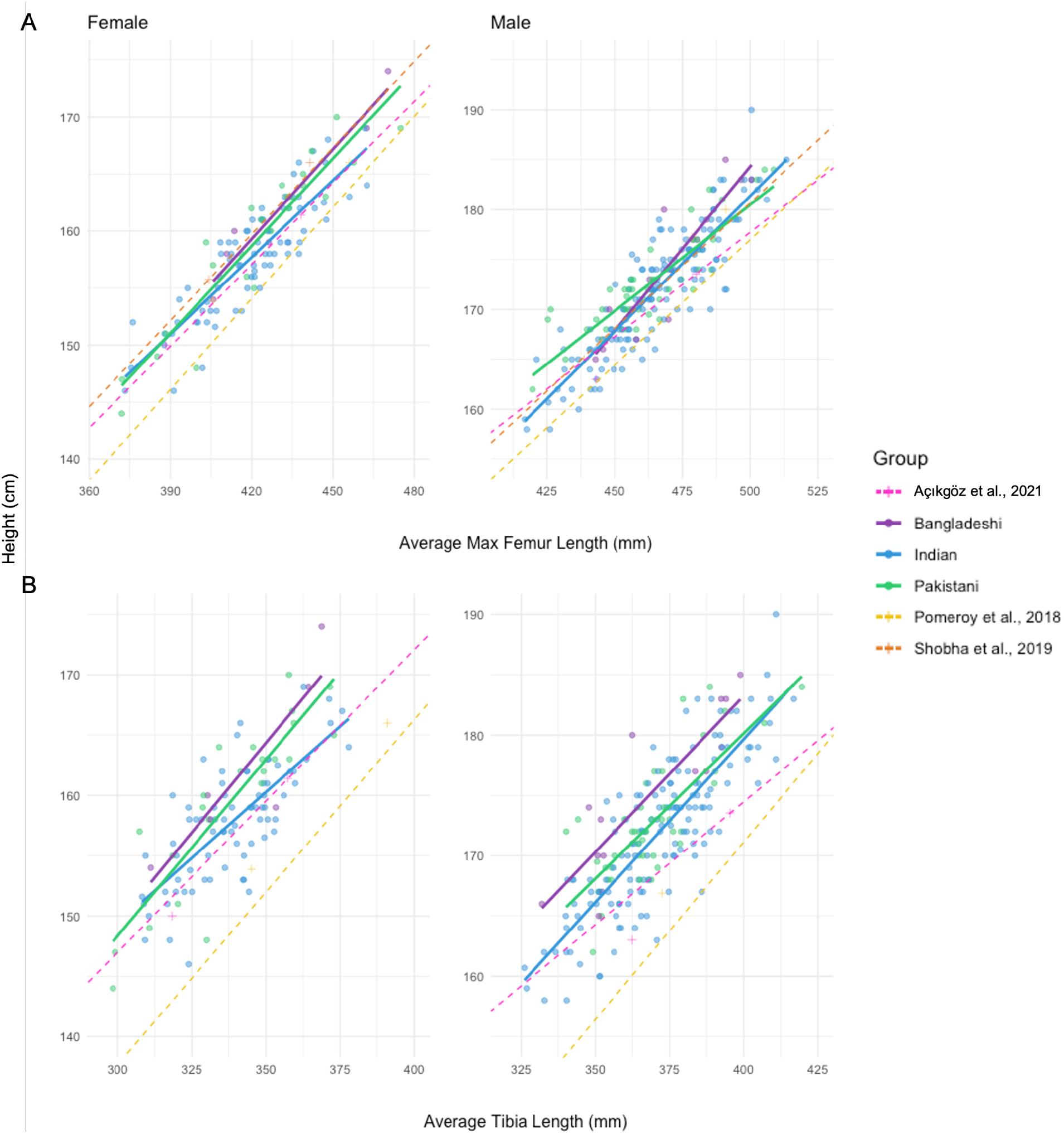
Univariate regression plots illustrating the relationships between stature and A) average maximum femoral lengths, and B) average tibiae lengths in the training sample and comparable groups (see publication details in Table 3). Circles and solid lines represent Bangladeshi (purple), Indian (blue), and Pakistani (green) standing height (cm) (UKB UDI 50-0.0) and automated skeletal measurements (mm). Dashed lines and + symbols represent the linear regression equations and descriptive statistics presented by Açikgöz et al., 2021 (pink), Pomeroy et al., 2018 (yellow), and Shobha et al., 2019 (orange). Only maximum femur length was reported by Shobha et al.

## Discussion

The results of the error assessment indicate that our proposed pipeline can produce accurate and automated inter-landmark skeletal measurements from DXA images. Additionally, it is promising that the test sample matched the training sample in landmark propagation acceptance rate (90%) despite demographic differences between the test, training, and atlas samples (Figure 1). No Bangladeshi participants were incorporated as atlas images, and the average stature across groups, samples, and sexes is not uniformly distributed (for example, see the female Bangladeshi and Pakistani heights in Figure 1). This suggests that the pipeline is robust to size and morphological variation between the atlas and moving images with regards to population-specific variation and participant stature. However, not all measurements performed equally well and several factors must be considered when implementing this automated phenotyping pipeline.

### Limitations

As with most data processing and analyses, the quality of the output is dependent on the quality of the input. Therefore, limitations were imposed by the quality and characteristics specific to the images passed through the pipeline. Within the UKB DXA imaging sample, participant body positioning is inconsistent. This affects automated skeletal phenotyping at two levels of the pipeline: initially, when deciding on landmark definitions, and later during image registration. Highly mobile regions (e.g. joints of the upper limb) are more difficult to uniformly define in 2D space, and this difficulty is compounded when standard anatomical positioning is not enforced. Despite prior efforts to standardize landmark placement at the proximal humerus, elbow, and wrist, the upper limb could not be incorporated into the current assessment as global body positioning was too variable for any of the image transformations to overcome during image registration (Cataldo-Ramirez et al., 2022). Additionally, many images did not include the elbow or forearm as these parts of the body were physically located outside of the scanning area. A global image registration approach may not be best suited for samples in which arm positioning is highly variable. Another fundamental consideration inherent to the imaging sample is the image resolution. DXA scans, while safe, convenient, and relatively cheap, are also less detailed than other forms of medical imaging. Some skeletal features and regions are simply not visible because of this (e.g., medial clavicles), and smaller skeletal measurements (e.g., acetabular breadth) are more influenced by image resolution than larger measurements (e.g., tibia length). This translates to a higher percent error for smaller measurements, as each pixel comprises a relatively larger proportion of the measurement. Automated smaller measurements may not be obtainable from whole body DXA scans, but it may be possible to extract them from regional DXA scans with smaller scales.

Other limitations were identified specific to the skeletal measurements characterized as “variable”. For trunk length and upper thoracic breadth, regional information is considered when manually placing the relevant landmarks (Table 2, Figure 3, SI Table 1) that the localized, nonlinear transformation step of the ANTs registration procedure can not incorporate. For example, vertebrae were counted and used as relative guidelines, and knowledge of skeletal variation was considered when manually placing rib and vertebral landmarks, but this information is not used by the ANTs transformation models. The other “variable” measurement, innominate height, was likely influenced by decisions enacted during image pre-processing. Because of the differences in opacity (and therefore, pixel density values) across skeletal regions, and the variability within UKB DXA image processing (e.g., visibility of soft tissue), some variation in image quality was still present after implementing the image pre-processing procedure. As a result, the pelvic regions needed to define innominate height were not always visible, likely contributing to the inconsistent performance of this automated skeletal measurement. Modifying the image normalization parameters to allow more soft tissue visibility may resolve this issue. Lastly, as inter- and intra-observer error of manual landmark placements were not tested, it is possible that landmark (im)precision also contributed to the inconsistent performance of the “variable” measurements.

### Utility

Results for the “high” performing measurements (pelvic breadths and lower limb lengths) suggest that they can be automatically extracted with acceptable precision. Comparisons with previous research on the utility of extracting linear skeletal metrics manually from DXA scans present technical error of measurement (TEM) (Pomeroy et al., 2018). Although our error assessment methods are not directly comparable to the TEM used in Pomeroy et al., the average percent errors for max femoral length (corresponding to 3.2 mm) and tibia length (corresponding to 4.1 mm) presented here fall below the interobserver TEM reported for manual femoral (6.1 - 6.6 mm) and tibial (5.3 - 5.7 mm) measurements. This indicates that the automated measurements from our pipeline, for which there are comparable, published error assessments, perform as well as or better than skeletal measurements manually obtained by independent observers.

### Recommendations

In addition to the pipeline presented here, quality control steps should also be implemented when reviewing the automated skeletal measurements, as no measurement reached a CCC value of 1. Checking for outliers and visualizing the associated propagated landmarks is recommended. Calculating basic descriptive statistics of the automated measurements is one way of easily identifying images with measurements outside of a +/- 2 or +/- 3 SD range. This can be performed independently on each measurement, or on proportional relationships between skeletal measurements of the same individual (e.g., outliers could be detected based on femoral max length and/or for femoral max length divided by stature). Depending on the size of the sample, images with poor propagations may be thrown out, manually landmarked, or additional steps may be required, such as repeating the pipeline incorporating more atlases or modifying the image pre-processing parameters.

## Conclusions

Bi-iliac breadth, pelvic aperture breadth, maximum femoral length, and tibia length can be reliably extracted from DXA scans by our automated phenotyping pipeline, allowing for researchers to leverage biobank-level databases. The skeletal measurements extracted with the pipeline presented here are consistent with previously published data reflecting body proportions of South Asian groups, providing realistic utility for research into skeletal health, secular changes and evolutionary trends in body proportions, and population-specific equations for stature estimation within forensic contexts. The performance of this pipeline also suggests that it can be scaled up to extract skeletal measurements from larger databases, specifically allowing for the power necessary to investigate phenotype-genotype relationships among stature, its components, and related skeletal traits.

Furthermore, by expanding the feasibility of phenotyping diverse populations, global references can be developed to combat the Eurocentricity in medical research and standards. This is a particularly important consideration, as AI and machine-learning methods become increasingly tied to biobank-level research. Machine learning algorithms are not immune to biases, and the repercussions of this within the context of medical research and healthcare interventions cannot be ignored (Kostick-Quenet et al., 2022; Obermeyer et al., 2019). Although the largest DXA imaging sample in the UK Biobank is composed of British, Irish, and/or “White” identifying participants, one of the goals of the study was to prioritize the inclusion of non-majority groups when developing this automated phenotyping pipeline. Future refinements of automated phenotyping of medical imaging data should purposefully mitigate the perpetuation of European biases in order to produce more inclusive and globally beneficial methods.

## Acknowledgements

The authors would like to thank the UK Biobank (Research Project Number 54084) and its participants, without whom this study, and others like it, would not have been possible; David Katz, Jay Devine, and Amanda Neves for their guidance and recommendations; Brenna M. Henn and Mark N. Grote for their invaluable input; and UC Davis for awarding C. Cataldo-Ramirez with the *Summer Graduate Student Researcher (GSR) Award for Engineering or Computer-related Applications and Methods*.

## References

Açikgöz, A., Erkman, A., Binokay, F., Göker, P., & Bozkir, M. (2021). Stature estimation from radiographic measurements in adult Anatolian population. International Journal of Morphology, 39(3).

Altman, D. G. (1991). Practical statistics for medical research Chapman and Hall. London and New York.

Asif, H., Alliey-Rodriguez, N., Keedy, S., Tamminga, C. A., Sweeney, J. A., Pearlson, G., … & Gershon, E. S. (2021). GWAS significance thresholds for deep phenotyping studies can depend upon minor allele frequencies and sample size. Molecular psychiatry, 26(6), 2048–2055.

Avants, B. B., Johnson, H. J., & Tustison, N. J. (2015). Neuroinformatics and the insight toolkit. Frontiers in Neuroinformatics, 9, 5.

Avants, B. B., Tustison, N., & Song, G. (2009). Advanced normalization tools (ANTS). Insight j, 2(365), 1–35.

Cataldo-Ramirez, C. C., Haddad, D., Amenta, N., & Weaver, T. D. (2022, April). Assessing the performance of Advanced Normalization Tools (ANTs) in automating skeletal phenotype extraction from large datasets of 2D images. Poster presented at the 91st Annual Meeting of the American Association of Biological Anthropologists, Denver, CO.

Dryden, I. L. & Mardia, K. V. (1998). Statistical shape analysis. John Wiley & Sons, New York.

Kassambara A, Mundt F (2020). _factoextra: Extract and Visualize the Results of Multivariate Data Analyses_. R package version 1.0.7, <https://CRAN.R-project.org/package=factoextra>.

Kostick-Quenet, K. M., Cohen, I. G., Gerke, S., Lo, B., Antaki, J., Movahedi, F., … & Blumenthal-Barby, J. S. (2022). Mitigating racial bias in machine learning. Journal of Law, Medicine & Ethics, 50(1), 92–100.

Lê, S., Josse, J., Husson, F. (2008). “FactoMineR: A Package for Multivariate Analysis.” Journal of Statistical Software, 25(1), 1–18. doi:10.18637/jss.v025.i01.

Li, Xiangrui (2022). xiangruili/dicm2nii (https://github.com/xiangruili/dicm2nii), GitHub. Retrieved September 5, 2022.

Njeh, C. F., Samat, S. B., Nightingale, A., McNeil, E. A., & Boivin, C. M. (1997). Radiation dose and in vitro precision in paediatric bone mineral density measurement using dual X-ray absorptiometry. The British journal of radiology, 70(835), 719–727.

Manurung, Auralius (2022). Elliptic fourier for shape analysis (https://www.mathworks.com/matlabcentral/fileexchange/32800-elliptic-fourier-for-shape-analysis), MATLAB Central File Exchange. Retrieved September 7, 2022.

MATLAB. (2022a). Natick, Massachusetts: The MathWorks Inc.

Obermeyer, Z., Powers, B., Vogeli, C., & Mullainathan, S. (2019). Dissecting racial bias in an algorithm used to manage the health of populations. Science, 366(6464), 447–453.

Pomeroy, E., Mushrif-Tripathy, V., Wells, J. C., Kulkarni, B., Kinra, S., & Stock, J. T. (2018). Stature estimation equations for South Asian skeletons based on DXA scans of contemporary adults. American journal of physical anthropology, 167(1), 20–31.

R Core Team (2022). R: A language and environment for statistical computing. R Foundation for Statistical Computing, Vienna, Austria. URL https://www.R-project.org/.

Shobha, Pravinkumar N. Kamaradgi, Pragnya Rao, Vijayakumar B Jatti. Estimation of stature from radiological length of femur among South - Indian adult population. International Journal of Contemporary Medical Research 2019;6(7):G1–G4

Yushkevich, P. A., Piven, J., Hazlett, H. C., Smith, R. G., Ho, S., Gee, J. C., & Gerig, G. (2006). User-guided 3D active contour segmentation of anatomical structures: significantly improved efficiency and reliability. Neuroimage, 31(3), 1116–1128.

